# Clearance of senescent decidual cells by uterine natural killer cells drives endometrial remodeling during the window of implantation

**DOI:** 10.1101/172593

**Authors:** Paul J Brighton, Yojiro Maruyama, Katherine Fishwick, Pavle Vrljicak, Shreeya Tewary, Risa Fujihara, Joanne Muter, Emma S Lucas, Taihei Yamada, Laura Woods, Raffaella Lucciola, Yie Hou Lee, Satoru Takeda, Sascha Ott, Myriam Hemberger, Siobhan Quenby, Jan J Brosens

**Affiliations:** Division of Biomedical Sciences, Clinical Science Research Laboratories, Warwick Medical School, University of Warwick, Coventry CV2 2DX, UK; Department of Obstetrics and Gynecology, Juntendo University Faculty of Medicine, Tokyo, 113-8421, Japan; Tommy’s National Centre for Miscarriage Research, University Hospitals Coventry & Warwickshire, Coventry CV2 2DX, UK; Centre for Trophoblast Research, University of Cambridge, Downing Street, Cambridge CB2 3EG, UK; Epigenetics Programme, The Babraham Institute, Babraham Research Campus, Cambridge CB22 3AT, UK; Obstetrics & Gynaecology-Academic Clinical Program, Duke-NUS Medical School, Singapore; KK Research Centre, KK Women’s and Children’s Hospital, Singapore

**Keywords:** endometrium, decidualization, senescence, uterine natural killer cells, homeostasis, implantation, senescence-associated secretory phenotype, IL-8, FOXO, granule exocytosis, senolytics

## Abstract

In cycling human endometrium, menstruation is followed by rapid estrogen-dependent growth. Upon ovulation, progesterone and rising cellular cAMP levels activate the transcription factor Forkhead box O1 (FOXO1) in endometrial stromal cells (EnSCs), leading to cell cycle exit and differentiation into decidual cells that control embryo implantation. Here we show that FOXO1 also causes acute senescence of a subpopulation of decidualizing EnSCs in an IL-8 dependent manner. Selective depletion or enrichment of this subpopulation revealed that decidual senescence drives the transient inflammatory response associated with endometrial receptivity. Further, senescent cells prevent differentiation of endometrial mesenchymal stem cells in decidualizing cultures. As the cycle progresses, IL-15 activated uterine natural killer (uNK) cells selectively target and clear senescent decidual cells through granule exocytosis. Our findings reveal that acute decidual senescence governs endometrial rejuvenation and remodeling at embryo implantation, and suggest a critical role for uNK cells in maintaining homeostasis in cycling endometrium.

## Introduction

Different mammalian species employ divergent strategies to ensure successful embryo implantation. In mice, synchronized implantation of multiple embryos (average 6-8) is dependent on a transient rise in circulating estradiol (E2) that not only renders the progesterone-primed endometrium receptive, but also activates dormant blastocysts for implantation (Paria et al., 1998). Upon breaching of the uterine luminal epithelium, implanting murine embryos trigger extensive remodeling of the endometrial stromal compartment. This process, termed decidualization, is characterized by local edema, influx of uNK cells and differentiation of stromal fibroblasts into specialized decidual cells that coordinate trophoblast invasion and placenta formation (Gellersen and Brosens, 2014; Zhang et al., 2013). Likewise, the human endometrium transiently expresses a receptive phenotype, lasting 2-4 days, during the mid-luteal phase of the cycle. However, this implantation window is not controlled by a nidatory E2 surge (de Ziegler et al., 1992; Groll et al., 2009), perhaps reflecting that synchronized implantation of multiple human embryos is neither required nor desirable. Further, decidualization of the stromal compartment is not dependent on an implanting embryo but initiated during the mid-luteal phase of each cycle in response to elevated circulating progesterone levels and increased intracellular cAMP production (Gellersen and Brosens, 2014). In parallel, CD56^bright^ CD16^−^ uNK cells accumulate in luteal phase endometrium. In pregnancy, uNK cells exert an evolutionarily conserved role in orchestrating vascular adaptation and trophoblast invasion (Hanna et al., 2006; Xiong et al., 2013), but their function in cycling human endometrium is unclear.

Differentiation of human endometrial stromal cells (EnSCs) into decidual cells is a multistep process (Gellersen and Brosens, 2014). Following cell cycle exit at G0/G1, decidualizing EnSCs first mount a transient pro-inflammatory response, characterized by a burst of free radical production and secretion of various chemokines and other inflammatory mediators (Al-Sabbagh et al., 2011; Lucas et al., 2016b; Salker et al., 2012). Exposure of the mouse uterus to this inflammatory secretome activates multiple receptivity genes, suggesting that the nidatory E2 surge in mice is supplanted by an endogenous inflammatory signal in the human uterus. Feedback loops purportedly limit the inflammatory decidual response to 2-4 days (Salker et al., 2012). The next decidual phase coincides with embedding of the implanted embryo into the stroma. At this stage, fully differentiated decidual cells, which are now tightly adherent and possess gap junctions (Laws et al., 2008), form an immune privileged matrix around the semi-allogenic conceptus (Erlebacher, 2013). In the absence of implantation, falling progesterone levels trigger a second inflammatory decidual response which, upon recruitment and activation of leukocytes, leads to tissue breakdown, focal bleeding and menstrual shedding of the superficial endometrial layer. Scar-free tissue repair involves activation of mesenchymal stem-like cells (MSCs) and epithelial progenitor cells that reside in the basal layer (Evans et al., 2016). Following menstruation, rising follicular E2 levels drive rapid tissue growth, which over ~10 days increases the thickness of the endometrium several-fold. Clinically, suboptimal endometrial growth is strongly associated with reproductive failure (Yuan et al., 2016); but how MSC activation followed by intense proliferation is linked to the decidual process is unclear.

FOXO1 is a core decidual transcription factor that controls cell cycle exit of EnSCs in response to differentiation signals and activates expression of decidual marker genes, such as *PRL* and *IGFBP1* (Park et al., 2016; Takano et al., 2007). Here we demonstrate that FOXO1 also induces acute senescence in a subpopulation of EnSCs. We show that the senescence-associated secretory phenotype (SASP) drives the initial auto-inflammatory decidual response linked to endometrial receptivity and provide evidence that uNK cells target and eliminate senescent decidual cells as the cycle progresses. Our findings reveal a hitherto unrecognized role for acute cellular senescence in endometrial remodeling at the time of embryo implantation; and suggest a major role for uNK cells in maintaining tissue homeostasis from cycle to cycle.

## Results

### Decidualization induces acute senescence in a subpopulation of EnSCs

To determine if cycling human endometrium harbor dynamic populations of senescent cells, we first stained primary EnSC cultures for senescence-associated β-galactosidase (SAβG) activity. At passage 1 (P1), SAβG^+^ cells were detectable in variable numbers in different cultures (Figure 1A). Strikingly, the number of SAβG^+^ cells increased markedly upon decidualization with 8-bromo-cAMP and medroxyprogesterone acetate (MPA, a progestin). Typically, SAβG^+^ cells formed islets surrounded by SAβG^-^ EnSCs in differentiating cultures (Figure 1A). Quantitative analysis confirmed a time-dependent increase in SAβG activity upon decidualization (Figure 1B). The abundance of SAβG^+^ cells in undifferentiated cultures declined upon passaging of cells (Figure S1A). Initially, this was paralleled by a reduction in SAβG activity, which was reversed at later passages (P6) (Figure S1B), presumably reflecting emerging replicative exhaustion of EnSCs (Figure S1C). However, even after ~60 days in continuous culture, exposure of EnSCs to a deciduogenic stimulus enhanced SAβG activity and triggered the appearance of SAβG^+^ cells (Figure S1A and S1B).

**Figure 1.**
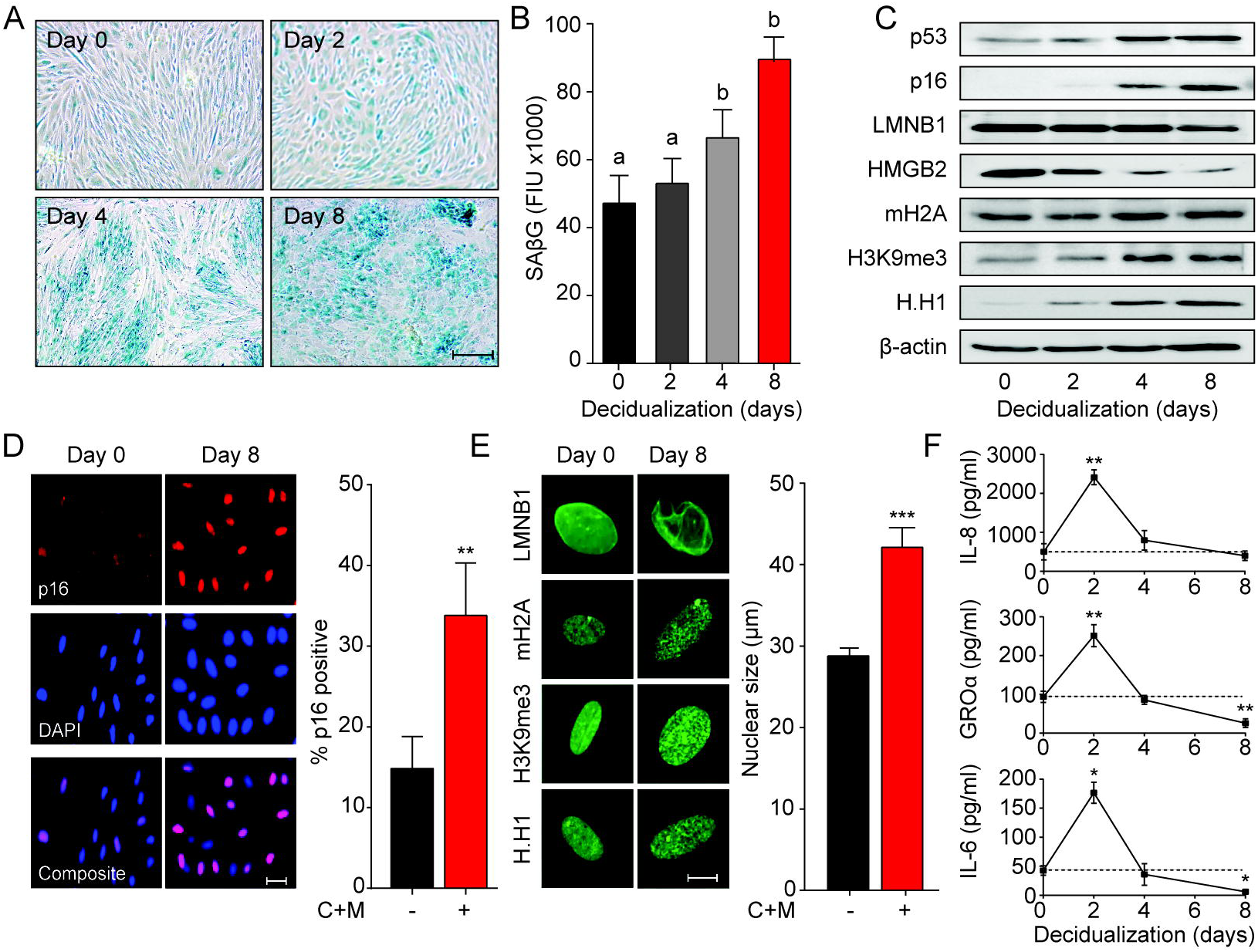
Decidualization induces acute senescence in a subpopulation of EnSCs. (A) Representative SAβG staining in undifferentiated EnSCs (Day 0) or cells decidualized for the indicated time points with 8-bromo-cAMP and MPA. Scale bar = 100 μm. (B) SAβG activity, expressed in fluorescence intensity units (FIU), in undifferentiated EnSCs (day 0) or cells decidualized for the indicated time points. (C) Representative Western blot analysis of p53, p16, LMNB1, HMGB2, mH2A, H3K9me3 and H.H1 levels in undifferentiated EnSCs and cells decidualized for the indicated time points. β-actin served as a loading control. (D) Left panel: representative immunofluorescence staining for p16 expression in undifferentiated cells and cells decidualized for 8 days. Nuclei were counterstained with DAPI. Scale bar = 50 μm. Right panel: percentage of p16^+^ cells. (E) Left panel: representative confocal microscopy images of undifferentiated (Day 0) or decidualized (Day 8) EnSCs immune-probed for LMNB1, mH2A, H3K9me3 and H.H1. Scale bar = 10 μm. Right panel: nuclear size of undifferentiated EnSCs (n = 48) and of cells first decidualized for 8 days with 8-br-cAMP and MPA (C+M) (n = 48) was measured in 3 primary cultures. (F) Secretion of IL-8, GROα, and IL-6 was measured in the supernatant of primary EnSCs collected every 48 h over an 8-day decidualization time-course. Data are mean ± SEM of 3 biological replicates unless stated otherwise. ^∗∗^ *P*< 0.01, ^∗∗∗^ *P*< 0.001. Different letters above the error bars indicate that those groups are significantly different from each other at *P* < 0.05. See also Figure S1.

Although SAβG activity is a commonly used biomarker for senescent cells, it lacks specificity (Matjusaitis et al., 2016). Hence, we examined the expression of other putative senescence markers in undifferentiated and decidualizing EnSCs. Many senescence signals converge onto the tumor suppressor protein p53 (p53) and induce the expression of cyclin-dependent kinase (CDK) inhibitors, leading to proliferative arrest and cell cycle exit (Munoz-Espin and Serrano, 2014; van Deursen, 2014). We reported previously that downregulation of MDM2 proto-oncogene, an E3 ubiquitin ligase, stabilizes p53 in differentiating EnSCs (Pohnke et al., 2004). Western blot analysis showed that p53 stabilization upon decidualization is paralleled by upregulation of p16^Ink4a^ (p16; Figure 1C), a CDK inhibitor presumed specific for senescence. Notably, confocal microscopy revealed that induction of p16 upon decidualization is confined to a subpopulation of EnSCs (Figure 1D). Loss of lamin B1 (LMNB1) and high mobility group box 2 (HMGB2) drives many of the chromatin and epigenetic changes that underpin cellular senescence (Aird et al., 2016; Sadaie et al., 2013). Decidualization resulted in downregulation of both effector proteins and a reciprocal increase in the histone H2A variant macroH2A (mH2A) and trimethylated lysine 9 on histone H3 (H3K9Me3) (Figures 1C and S1E). This histone variant and modification are involved in senescence-associated heterochromatin formation (SAHF). Unexpectedly, the nucleosome linker histone H1 (H.H1), which purportedly is lost in senescent cells (Funayama et al., 2006), was upregulated upon decidualization (Figures 1C and S1E). These observations were confirmed by immunofluorescence confocal microscopy (Figure 1E, left panel), which also revealed that the nuclei of EnSC increase in size (~ 40%) upon decidualization (Figure 1E, right panel). Next, we examined if the transient inflammatory decidual response also encompasses secreted factors typical of the canonical senescence associated secretory phenotype (SASP). As shown in Figure 1F, secretion of IL-8 (CXCL8), GROα (CXCL1), and IL-6 peaked transiently on day 2 of decidualization and returned to baseline by day 4. By day 8, the level of secretion of GROα and IL-6 was lower than that observed in undifferentiated cultures.

Taken together, the data reveal striking similarities between cellular senescence and differentiation of EnSCs into decidual cells. However, only a subpopulation of EnSCs became strongly SAβG^+^ or expressed p16 upon decidualization. Further, while SASP is often a sustained response, decidual inflammation is temporally restricted to 2-4 days.

### Temporal regulation of senescent cell populations in cycling endometrium

To extrapolate these observations to the *in vivo* situation, protein lysates from whole endometrial biopsies were subjected to Western blot analysis. As the cycle progresses from the proliferative to the secretory phase, the abundance of p53, p16, LMNB1, HMBG2, mH2A and H3K9me3 in the endometrium mimicked the changes observed in decidualizing EnSC cultures (Figure 2A). Further, analysis of snap-frozen biopsies showed a sharp increase in SAβG activity upon transition from proliferative to early-secretory endometrium with levels peaking in the late-luteal phase (Figure 2B). Disintegration of the stromal compartment upon cryosectioning of frozen tissue samples precluded a meaningful analysis of SAβG^+^ cells. Instead, we used immunohistochemistry to examine the abundance and tissue distribution of p16^+^ cells in 308 formalin-fixed endometrial biopsies obtained during the peri-implantation window, i.e. 6 to 12 days after the luteinizing hormone (LH) surge (Figure 2C). The statistical distribution of p16^+^ cells in the glandular epithelium, luminal epithelium and stromal compartment is presented as a centile graph (Figure 2D). Interesting, p16^+^ cells were most prevalent in both the glandular and luminal epithelium at LH+10 and +11, which coincides with the onset of the late-luteal phase of the cycle. The relative abundance of p16^+^ cells was ~10-fold higher in the luminal compared to the glandular compartment. Typically, stretches of p16^+^ cells were interspersed by p16^-^ cells in the luminal epithelium (Figure 2C). By contrast, p16^+^ cells gradually increased in the stromal compartment during the mid-luteal phase and this was accelerated in late-luteal endometrium. Occasionally, swirls of p16^+^ cells were observed in the stroma, seemingly connecting the deeper regions of the endometrium to p16^+^ luminal epithelial cells (Figure 2C). Collectively, the data indicate that the endometrium harbors dynamic and probably spatially organized populations of senescent cells during the luteal phase of the cycle.

**Figure 2.**
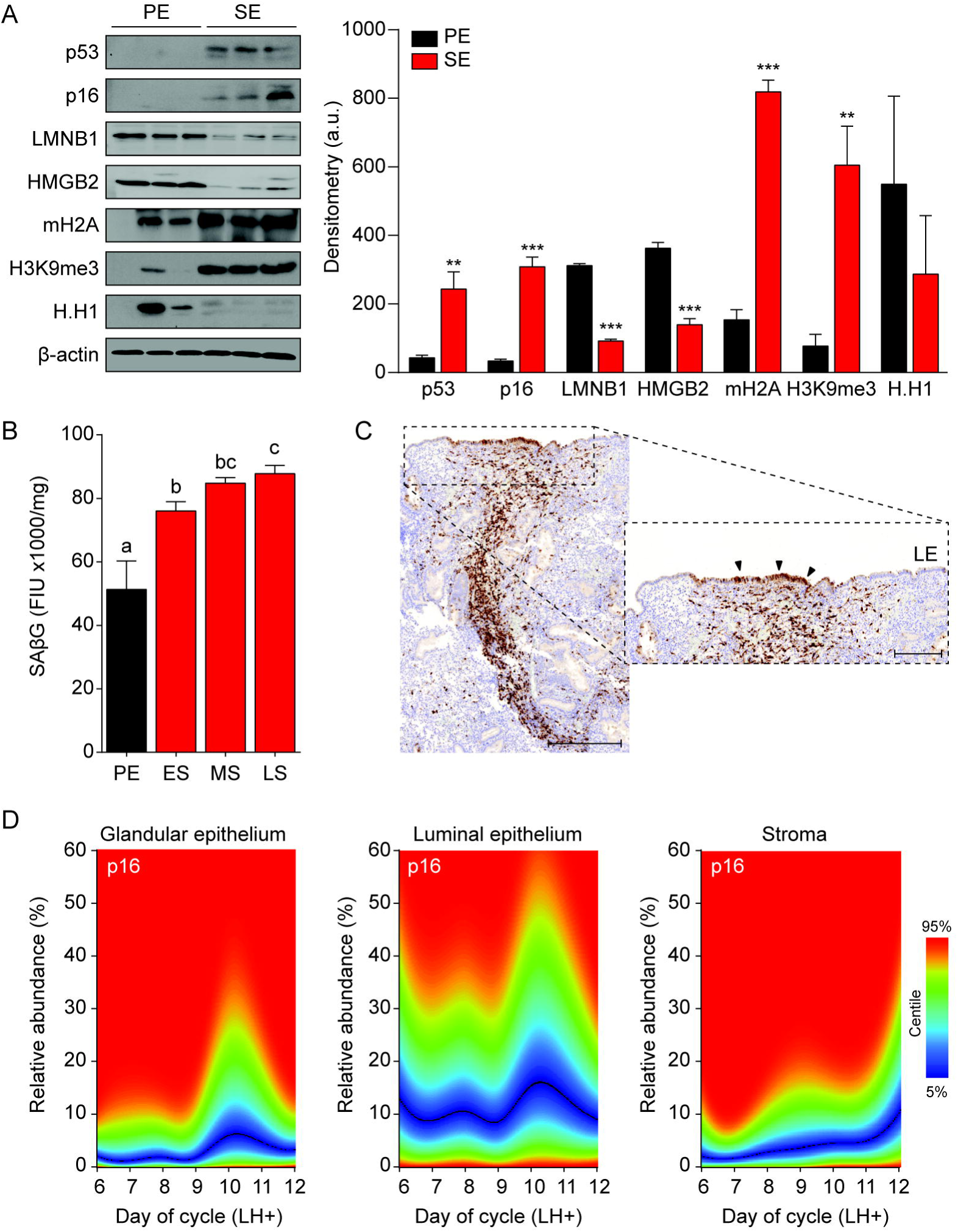
Senescent cells in cycling human endometrium. (A) Left panel: representative Western blot analysis of p53, p16, LMNB1, HMGB2, mH2A, H3K9me3 and H.H1 levels in whole tissue biopsies from proliferative endometrium (PE) and secretory endometrium (SE). β-actin served as a loading control. Right panel: protein levels quantified relative to β-actin by densitometry and expressed as arbitrary units (a.u.). (B) SAβG activity, expressed in fluorescence intensity units (FIU) / mg protein, was measured in biopsies from proliferative endometrium (PE; n = 7), early-secretory (ES; n = 9), midsecretory (MS; n = 38) and late-secretory (LS; n = 19) endometrium. (C) Immunohistochemistry demonstrating distribution of p16^+^ cells in the stromal compartment and luminal epithelium. Scale bars = 200 μm (D) The abundance of p16^+^ cells during the luteal phase in glandular epithelium, luminal epithelium and stroma compartment was analyzed by color deconvolution using ImageJ software in 308 LH-timed endometrial biopsies (average 48 samples per time point; range: 22 to 69). The centile graphs depict the distribution of p16^+^ cells across the peri-implantation window in each cellular compartment. Color key is on the right. Data are mean ± SEM of 3 biological replicates unless stated otherwise. ^∗∗^ *P*< 0.01, ^∗∗∗^ *P*< 0.001. Different letters above the error bars indicate that those groups are significantly different from each other at *P* < 0.05.

### FOXO1 drives EnSC differentiation and senescence

To gain insight into the mechanism that drives decidual senescence, SAβG activity was measured in cultured EnSCs treated for 8 days with either 8-bromo-cAMP, MPA or a combination. Both 8-bromo-cAMP and MPA were required for significant induction of SAβG activity (*P* < 0.05) (Figure 3A). In differentiating EnSCs, cAMP and progesterone signaling converge on FOXO1, a core decidual transcription factor responsible for cell cycle arrest and induction of decidual marker genes, such as *PRL* and *IGFBP1* (Takano et al., 2007). Interestingly, FOXO1 was also shown to induce cellular senescence of ovarian cancer cells treated with progesterone (Diep et al., 2013). siRNA-mediated knockdown of FOXO1 in EnSCs not only abolished the induction of *PRL* and *IGFBP1* (Figure S2A) but also the surge in IL-8, GROα, and IL-6 secretion upon treatment of cultures with 8-bromo-cAMP and MPA (Figure 3B). After 8 days of decidualization, FOXO1 knockdown was less efficient but nevertheless sufficient to significantly blunt SAβG activity (*P* < 0.05) (Figure 3C). Several components of the SASP have been implicated in autocrine/paracrine propagation of senescence, including IL-8 acting on CXCR2 (IL-8 receptor type B) (Acosta et al., 2008). In agreement, decidualization of EnSCs in the presence of SB265610, a potent CXCR2 inhibitor, attenuated SAβG activity in a dose-dependent manner (Figure 3D). Conversely, recombinant IL-8 upregulated SAβG activity in undifferentiated EnSCs in a concentration-dependent manner (Figure 3D), although spatial organization of SAβG^+^ cells into ‘islets’ was not observed (Figure S2B). Further, siRNA-mediated *CXCL8* (coding IL-8) knockdown in undifferentiated EnSCs not only blunted the surge in IL-8 secretion upon decidualization (Figure S2B), but also abolished the increase in SAβG activity (Figure 3E). Unexpectedly, IL-8 knockdown compromised the induction of *PRL* and *IGFBP1* in cultures treated with 8-bromo-cAMP and MPA (Figure 3F), indicating that autocrine/paracrine signals involved in EnSC differentiation also drive decidual senescence.

**Figure 3.**
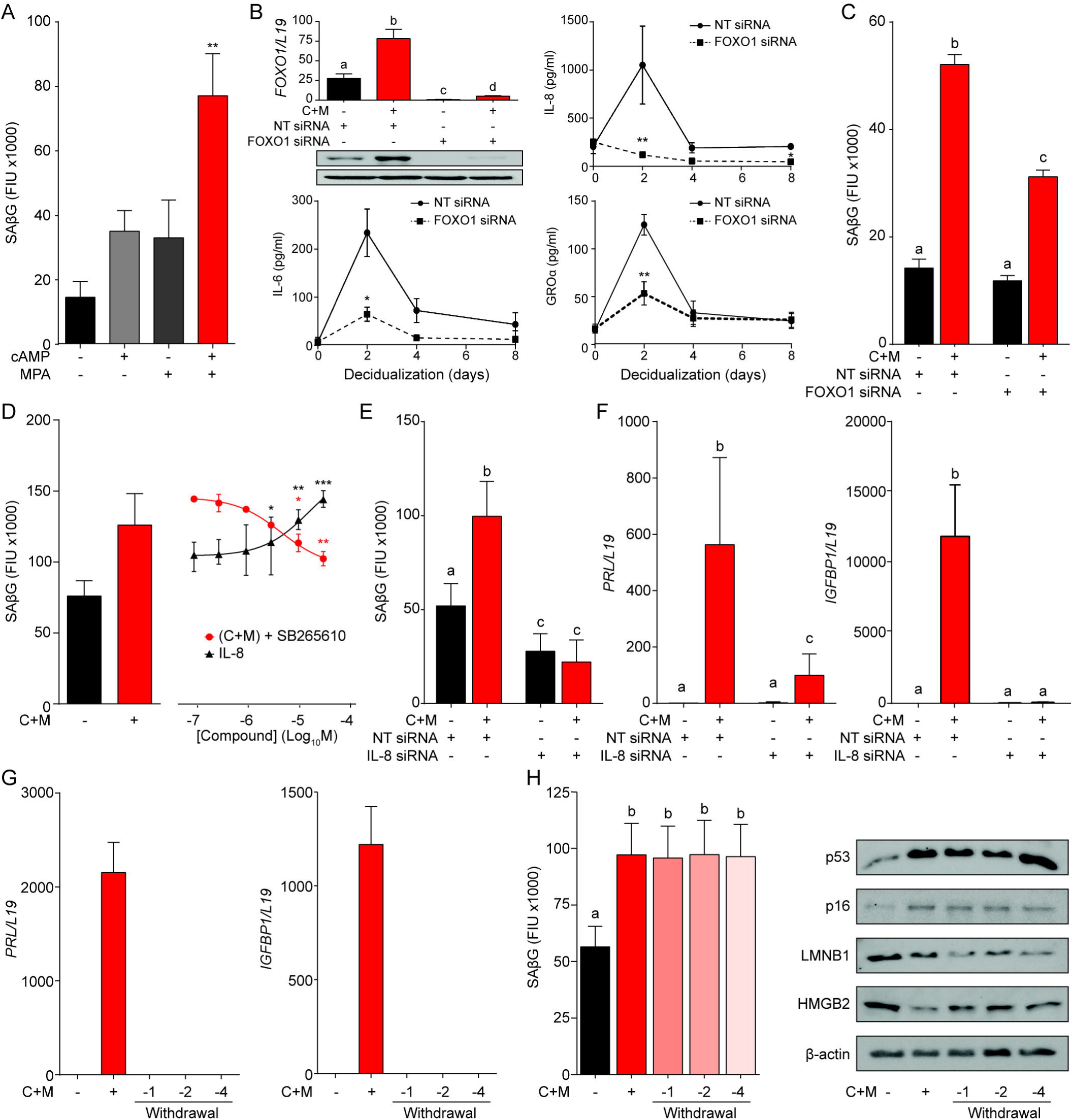
A FOXO1 / IL-8 axis drives EnSC differentiation and senescence. (A) SAβG activity in EnSCs either undifferentiated, or decidualized for 8 days with 8-bromo-cAMP, MPA, or a combination. (B) Top left panel: *FOXO1* mRNA levels in undifferentiated EnSCs and cells treated with 8-br-cAMP and MPA (C+M) following transfection with non-targeting (NT) or FOXO1 siRNA. Other panels: Secretion of IL-8, IL-6 and GROα was measured following FOXO1 knockdown in the supernatant of primary EnSCs every 48 h over an 8-day decidualization time-course. (C) SAβG activity in EnSCs following transfection with NT or FOXO1 siRNA. The cultures either remain untreated or decidualized for 8 days. (D) SAβG activity in undifferentiated EnSCs treated for 8 days with increasing concentrations of recombinant IL-8 and in cells decidualized for 8 days in the presence of increasing concentrations of the CXCR2 antagonist, SB265610. (E) SAβG activity in EnSCs following transfection with IL-8 siRNA. The cultures either remain untreated or decidualized for 8 days. (F) *PRL* and *IGFBP1* transcript levels in EnSCs following transfection with IL-8 siRNA. The cultures either remain untreated or decidualized for 8 days. (G) *PRL* and *IGFBP1* expression in undifferentiated EnSCs, cells decidualized for 8 days, and upon withdrawal of 8-br-cAMP and MPA (C+M) for the indicated days. (H) Left panel: SAβG activity in undifferentiated EnSCs, cells decidualized for 8 days, and following withdrawal of C+M for the indicated days. Right panel: representative Western blot analysis of p53, p16, LMNB1 and HMGB2 levels in undifferentiated EnSCs, cells decidualized for 8 days, and following withdrawal of C+M for the indicated days. β-actin served as a loading control. Data are mean ± SEM of 3 biological replicates unless stated otherwise. ^∗^ *P*< 0.05, ^∗∗^ *P*< 0.01 and ^∗∗∗^ *P*< 0.005. Different letters above the error bars indicate that those groups are significantly different from each other at *P* < 0.05. See also Figure S2.

In an attempt to block decidual senescence selectively, EnSCs were differentiated in the presence or absence of the mTOR inhibitor rapamycin, a pharmacological repressor of replicative senescence (Demidenko et al., 2009). Rapamycin prevented expansion of SAβG^+^ cells upon decidualization but expression of *PRL* and *IGFBP1* was again compromised (Figures S2C and S2D). By contrast, withdrawal of 8-bromo-cAMP and MPA from cultures first decidualized for 8 days reversed the induction of decidual marker genes (Figure 3G), albeit without impacting on either SAβG activity or expression of p53, p16, LMNB1 and HMGB1 (Figure 3H). Taken together, the data demonstrate that FOXO1 drives both differentiation and senescence of distinct EnSC subpopulations in an IL-8 dependent manner; however, while expression of differentiation markers requires continuous cAMP and progestin signaling, the senescent phenotype does not.

### Pleiotropic functions of senescent decidual cells

The abundance of SAβG^+^ cells in undifferentiated cultures correlated closely with the number of SAβG^+^ cells upon decidualization (Pearson’s *r* = 0.97, *P* < 0.0001) (Figure S3A). A congruent correlation was apparent upon measuring SAβG activity in paired undifferentiated and decidualizing cultures (Pearson’s *r* = 0.91, *P* < 0.0001) (Figure 4A); inferring that senescent decidual cells arise from stressed (presenescent) EnSCs. Hence, we tested if decidualization-associated senescence could be blocked by pretreating undifferentiated cultures with senolytic drugs. Exposure of primary EnSCs for 72 h to increasing concentrations of ABT-263 (Navitoclax), a BCL-XL inhibitor (Zhu et al., 2016), had no discernible impact on the induction of SAβG activity upon decidualization (Figure S3B). By contrast, pretreatment of primary cultures with dasatinib, a broad-spectrum tyrosine kinase inhibitor (Childs et al., 2017; Zhu et al., 2015), inhibited SAβG activity upon decidualization in a dose-dependent manner (Figure S3C). Conversely, to increase the abundance SAβG^+^ cells, undifferentiated EnSCs were treated with the CDK4/CDK6 inhibitor palbociclib (PD0332991), a functional p16 mimetic (Mosteiro et al., 2016). Dose-response analyses showed that treatment with palbociclib for 4 days was sufficient to increase SAβG activity in undifferentiated EnSCs to the level observed in cells decidualized with 8-bromo-cAMP and MPA for 8 days (Figure S3C). Notably, neither dasatinib nor palbociclib pretreatment impacted on the induction of *PRL* or *IGFBP1* upon decidualization (Figure 4C). However, dasatinib pretreatment markedly blunted the surge in IL-8, IL-6, and GROα secretion that characterizes the initial decidual phase. By contrast, this auto-inflammatory decidual response was amplified in response to palbociclib pretreatment (Figure 4D).

**Figure 4.**
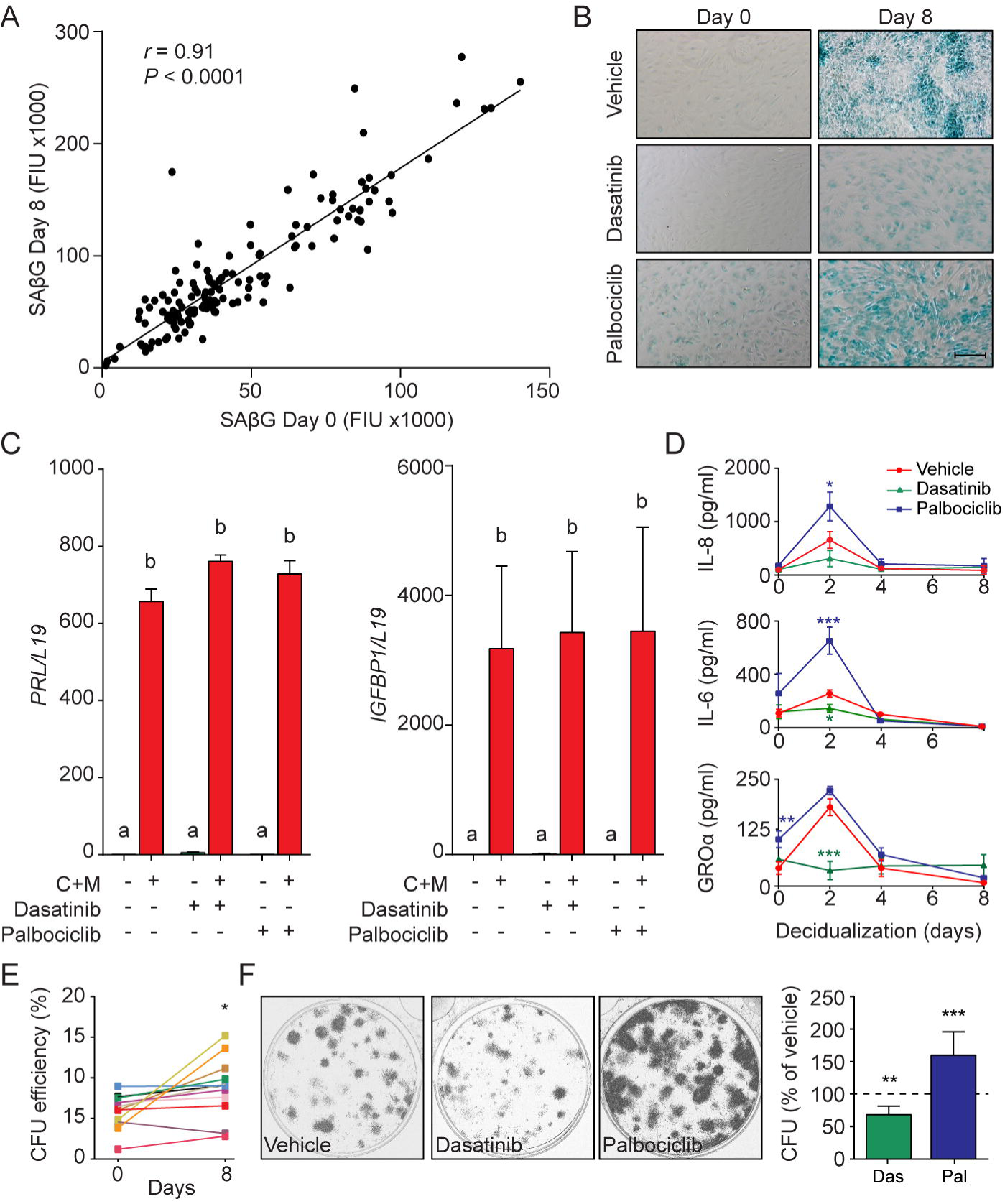
functions of senescent decidual cells. (A) Pearson’s correlation analysis of SAβG activity in 75 matched undifferentiated primary cultures and cultures decidualized for 8 days. (B) Representative SAβG staining in undifferentiated (Day 0) and decidualizing EnSCs (Day 8) following 4 days of pretreatment with vehicle, dasatinib (250 nM) or palbociclib (1 μM). Scale bar = 100 μm. (C) *PRL* and *IGFBP1* mRNA expression in response to pretreatment with vehicle, dasatinib or palbociclib. The cultures then remained undifferentiated or were decidualized for 8 days. (D) IL-8, IL-6 and GROα secretion was measured every 48 h in the supernatant of primary EnSCs decidualized for the indicated time-points following pretreatment with vehicle, dasatinib or palbociclib. (E) Colony forming unit (CFU) activity in paired EnSC cultures that either remain undifferentiated (Day 0) or were decidualized for 8 days (n = 10). (F) Left panel: representative clonogenic assays established from EnSC cultures first pretreated with vehicle, dasatinib or palbociclib and then decidualized for 8 days. Right panel: CFU activity in EnSC cultures first pretreated with vehicle, dasatinib or palbociclib and then decidualized for 8 days. Data are mean ± SEM of 3 biological replicates unless stated otherwise. ^∗^ *P*< 0.05, ^∗∗^ *P*< 0.01 and ^∗∗∗^ *P*< 0.001. Different letters above the error bars indicate that those groups are significantly different from each other at *P* < 0.05. See also Figure S3.

In other cell systems, transient - but not prolonged - exposure to SASP has been shown to promote tissue rejuvenation by reprogramming committed cells into stem-like cells (Ritschka et al., 2017). We reasoned that SASP-dependent tissue rejuvenation during the window of implantation may be relevant for the transition of the cycling endometrium into the decidua of pregnancy. To test this hypothesis, we examined the clonogenic capacity of paired undifferentiated cells and cells decidualized for 8 days. Analysis of 12 primary cultures demonstrated decidualization is associated with a modest but significant increase in colony-forming cells (*P* < 0.05), although the response varied between primary cultures (Figure 4E). However, pretreatment of undifferentiated cultures with dasatinib or palbociclib consistently increased and decreased the clonogenic capacity of decidualizing cultures, respectively (Figure 4F). Likewise, rapamycin also depleted decidualizing EnSC cultures of clonogenic cells (Figure S3D). Taken together, the data suggest that senescent decidual cells produce a transient inflammatory environment that not only renders the endometrium receptive but also increases tissue plasticity prior to pregnancy.

### Immune clearance of senescent decidual cells

Recognition and elimination of senescent cells by immune cells, especially NK cells, play a pivotal role in tissue repair and homeostasis (Iannello and Raulet, 2013; Krizhanovsky et al., 2008). During the luteal phase, uNK cells, characterized by their CD56^bright^ cell surface phenotype (Figure S4A), are by far the dominant endometrial leukocyte population. Analysis of a large number of LH-timed endometrial biopsies (n = 1,997) demonstrated that the abundance of CD56^+^ uNK cells in the endometrial stromal compartment increases on average 3-fold between LH+5 and +12; although inter-patient variability was marked (Figure 5A, left panel). The heatmap in Figure 5A (right panel) depicts the uNK cell centiles across the peri-implantation window. Notably, uNK cells often appear to amass in edematous areas that are relatively depleted of stromal cells, especially during the transition from the mid- to late-luteal phase of the cycle (Figure 5B, left panel). Quantitative analysis of 20 biopsies obtained between LH+9 and LH+11 confirmed the inverse correlation between the density of uNK cells and endometrial stromal cells (Figure 5B, right panel). In co-culture, uNK cells isolated from secretory endometrium had no impact on proliferation or viability of undifferentiated EnSCs (Figure S4B, left panel). By contrast, co-culture of uNK cells with EnSCs first decidualized for 8 days resulted in loss of cell viability. Visually, uNK cells transformed the monolayer of decidual cells into a honeycomb pattern with cell-free islets (Figure 5C).

**Figure 5.**
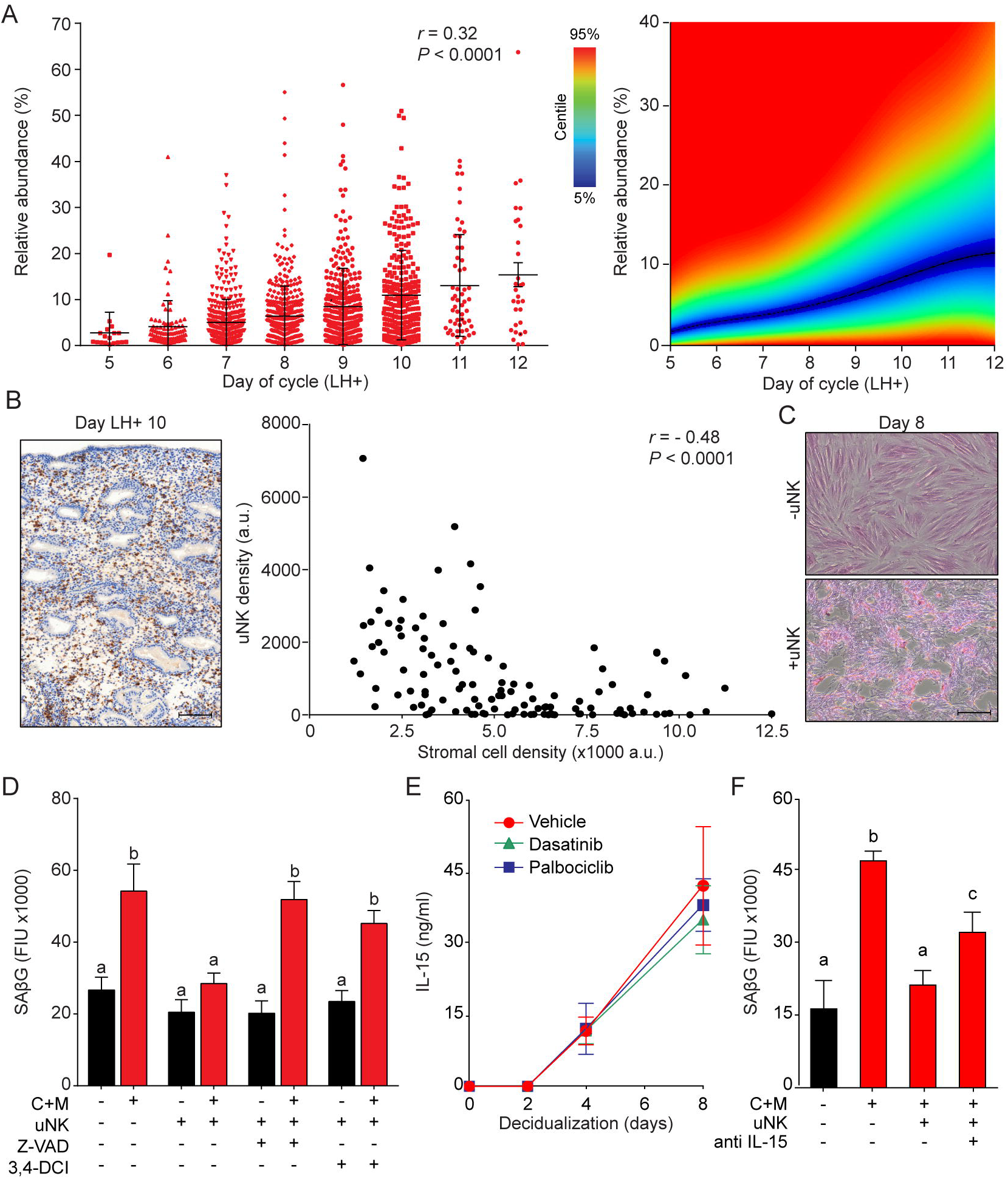
uNK cell mediated immune surveillance and clearance of senescent cells. (A) Left panel: uNK cell density in the subluminal stroma was quantified using a standardized immunohistochemistry protocol in LH timed endometrial biopsies (n = 1,997). Right panel: corresponding centile graph. Color code on the left. (B) Left panel: example of the tissue distribution of CD56^+^ uNK cells (brown staining) at LH+10. Scale bar = 250 μm. Right panel: Pearson’s correlation analysis of stromal cell and uNK cell densities. A total of 80 randomly selected images from 20 biopsies were analyzed. (C) Representative images of an eosin stained primary culture decidualized for 8 days incubated for 18 h with or without uNK cells isolated from luteal phase endometrium. Scale bar = 100 μm. (D) SAβG activity in undifferentitated or day 8 decidualized EnSCs co-cultured with or without uNK cells in the presence or absence of the apoptosis inhibitor Z-VAD-FMK (Z-VAD, 10 μM) or the granzyme activity inhibitor 3,4-DCI (25 μM). (E) Secretion of IL-15 secretion was measured every 48 h in the supernatant of primary EnSCs decidualized for the indicated time-points following pretreatment with vehicle, dasatinib (250 nM) or palbociclib (1 μM). (F) SAβG activity in undifferentitated or day 8 decidualized EnSCs co-cultured with or without uNK cells in the presence or absence of an IL-15 blocking antibody (1μg/ml). Data are mean ± SEM of 3 biological replicates unless stated otherwise. Different letters above the error bars indicate that those groups are significantly different from each other at *P* < 0.05. See also Figure S4.

These observations suggested that uNK cells actively eliminate senescent EnSCs but only upon decidualization. In agreement, co-culture of EnSCs with uNK eliminated the induction of SAβG activity upon decidualization without affecting basal activity in undifferentiated cells (Figure 5D). Two independent mechanisms underpin NK cell-mediated clearance of stressed cells (Sagiv et al., 2013). First, binding of the NK cell surface ligands TRAIL and FAS ligand (FasL) to the corresponding receptors on target cells can lead to caspase activation and cell death. However, incubation of primary EnSCs with increasing concentrations of FasL or TRAIL had no impact on SAβG activity in either undifferentiated or decidualizing cells (Figure S4C), inferring that death receptor activation in uNK cells is not required for senolysis. The second mechanism involves secretion by activated NK cells of granules containing perforin and granzyme (A, B). Perforin forms pores in the plasma membrane of target cells and triggers apoptosis upon release of granzyme into the cytoplasm (Chowdhury and Lieberman, 2008). As shown in Figures 5D and S4B, both the pan-caspase inhibitor Z-VAD-FMK and the granzyme B inhibitor 3,4-Dichloroisocoumarin (3,4-DCI) negated the impact of uNK cells on SAβG activity and cell viability in decidualizing cultures. To explore why uNK cell-mediated clearance of SAβG^+^ EnSCs is restricted to decidualizing cultures, we focused on IL-15, a pivotal cytokine that regulates NK cell proliferation and activation (Marcais et al., 2014). IL-15 secretion was below the level of detection in undifferentiated cells but, after a lag-period of 2 days, rose markedly upon decidualization of EnSCs in a time-dependent manner (Figure 5E). Notably, pretreatment of cultures with dasatinib or palbociclib had no impact on impact on IL-15 secretion, suggesting that decidual cells orchestrate the uNK-mediated clearance of their senescent counterparts (Figure 5E). Incubation of co-cultures with an IL-15 blocking antibody antagonized, at least partly, uNK cell-mediated clearance of senescent decidual cells (Figure 5F).

### Tissue homeostasis

Our findings indicate that endometrial homeostasis during the luteal phase is dependent on balancing induction and clearance of senescent decidual cells. We speculated that this process is *a priori* dynamic, which should be reflected in varying numbers of uNK cells in different cycles. As proof of concept, we quantified uNK cells in biopsies from 3 patients obtained around the same time in the mid-luteal phase (± 1 day) in 3 different cycles. As shown in Figure 6A, the abundance of uNK cells in the subluminal endometrial stroma can vary profoundly between cycles. As levels both rose and fell, the observed inter-cycle changes in uNK cell density are unlikely triggered by the tissue injury caused by the biopsy, although an impact on the magnitude of change cannot be excluded. Additional examples of uNK cell fluctuations in two consecutive cycles are presented in Figure S5A.

**Figure 6.**
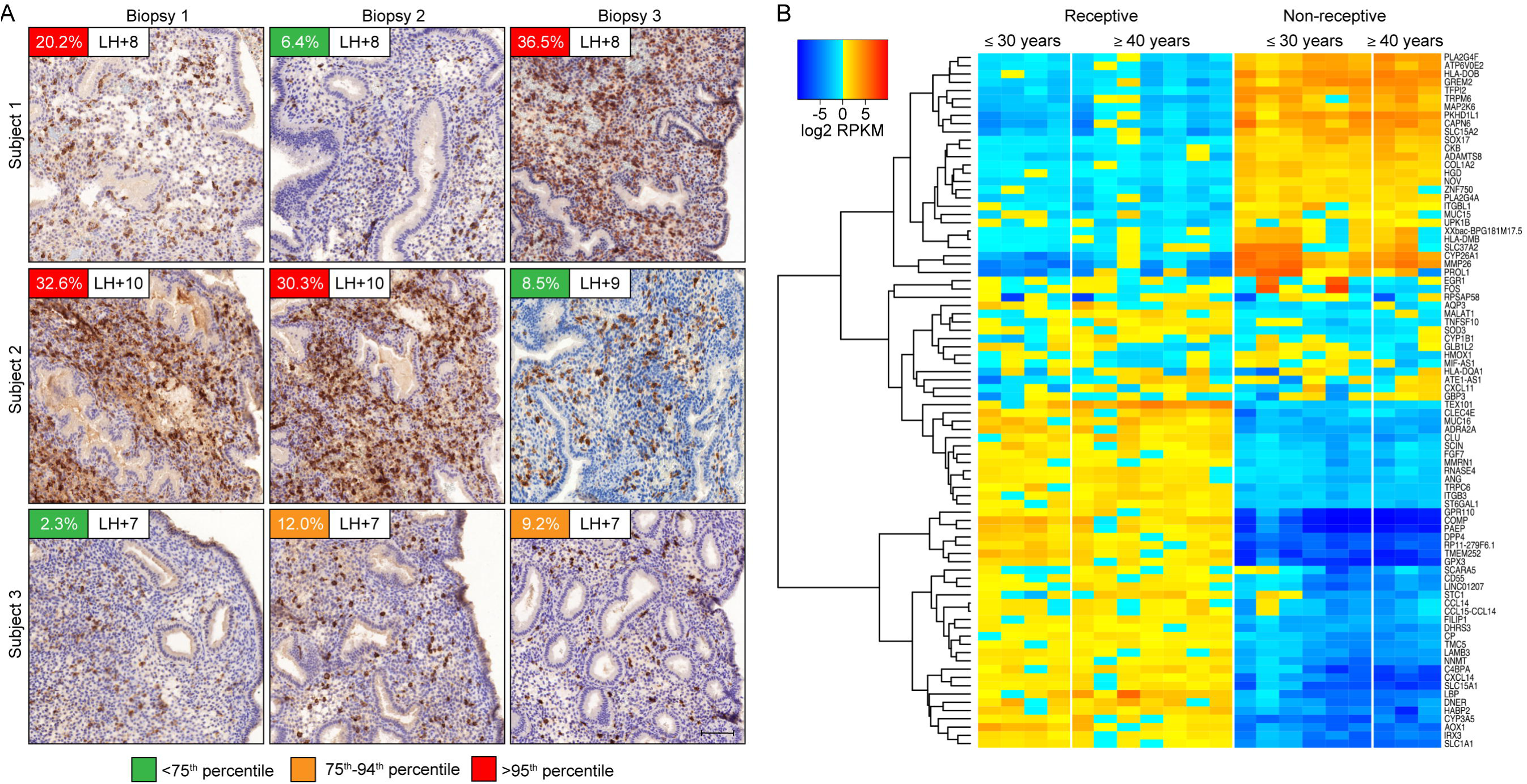
(A) CD56 immunohistochemistry of LH-timed endometrial biopsies obtained in 3 different cycles in 3 subjects. The day of the biopsy and the percentage of CD56^+^ uNK cells versus stromal cells are indicated. The color of the box indicates the percentile range of uNK when adjusted for the day of biopsy. Scale = 200 μm. (B). Heatmap showing that the 84 differentially expressed genes identified by intensity difference analysis (*P* < 0.05) following RNA-sequencing of endometrial biopsies of women aged ≤ 30 years or ≥ 40 years are accounted for by the receptive status of the biopsy. Note that more biopsies from the older group expressed a receptive phenotype when compared to samples from younger women. See also Figure S5.

Cyclic surveillance and elimination of senescent cells should protect the endometrium against chronological ageing. To substantiate this hypothesis, we performed RNA-sequencing on LH-timed endometrium biopsies obtained from 10 women aged ≤ 30 years and 10 women aged ≥ 40 years (Gene Expression Omnibus accession no. GSE102131). The samples were matched for body mass index, parity and day of biopsy but were separated by approximately ~170 menstrual cycles (Table S1). A total of 84 genes were identified as differentially expressed between the two groups (Figure S5B). However, 7 biopsies in the older age group expressed a receptive phenotype compared to 4 samples from younger women (Figure 6B). Thus, differential gene expression was accounted for by the state of receptivity of individual biopsies but not age. Taken together, the data suggest that cyclic endometrial senescence and rejuvenation may lead to short-term fluctuations in endometrial homeostasis during the window of implantation but long-term functional stability.

## Discussion

In contrast to chronic senescence associated with organismal ageing, acute senescence is a tightly orchestrated biological process implicated in embryo development, wound healing and tissue repair. Typically, acute senescent cells produce a context-specific SASP with defined paracrine functions and self-organize their elimination by various immune cells (van Deursen, 2014). Here we provide evidence that acute decidual senescence is a pivotal process that coordinates acquisition of a receptive phenotype with endometrial remodeling and rejuvenation during the implantation process. We reported previously that decidual transformation of primary EnSCs is a stepwise process that starts with a NOX4-dependent burst of free radicals and release of multiple inflammatory mediators (Al-Sabbagh et al., 2011; Lucas et al., 2016b; Salker et al., 2012). Exposure of the mouse uterus to this inflammatory secretome induced expression of multiple implantation genes and enabled efficient implantation of *in vitro* cultured mouse embryos (Salker et al., 2012). We now demonstrate that this nidatory decidual signal is driven foremost by acute senescence of a subpopulation of EnSCs. The close correlation between SAβG activity before and after decidualization suggests that polarization of EnSCs upon cell cycle exit into differentiating and senescent cells is not stochastic but determined by the level of replicative stress incurred by individual EnSCs during the preceding proliferative phase. Acute senescence rejuvenates the receptive endometrium through two distinct mechanisms. First, decidual SASP not only ‘locks in’ endometrial MSCs upon decidualization but, dependent on the amplitude of the inflammatory response, also de-differentiates more committed cells into clonogenic MSCs. Arguably, an adequate MSC population may be essential for expansion of the decidua in pregnancy. Second, clearance of senescent decidual cells upon uNK cell activation ensures that the embryo embeds in a preponderance of mature decidual cells. In co-culture, uNK cell mediated clearance of SAβG^+^ cells transformed the decidual cell monolayer into a honeycomb pattern. If recapitulated *in vivo*, this observation suggests a role for uNK cells in creating ingresses in the tightly adherent decidual cell matrix to facilitate trophoblast invasion and anchoring of the conceptus. Compared to undifferentiated EnSCs, decidual cells are highly resistant to various stress signals, convert inactive cortisone into cortisol through the induction of 11β-hydroxysteroid dehydrogenase type 1, and protect the embryo-maternal interface from influx of T-cells by silencing genes coding for key chemokines (Erlebacher, 2013; Gellersen and Brosens, 2014). Taken together, these observations suggest that senescent decidual cells trigger a dynamic tissue reaction that ultimately results in enclosure of the conceptus into an immune-privileged decidual matrix. In pregnancy, uNK cells express senescence markers and are proangiogenic rather than cytotoxic (Rajagopalan and Long, 2012). Whether prior exposure to senescent decidual cells contributes to this gestational phenotype of uNK cells is an intriguing but as yet untested possibility.

Notably, p16^+^ epithelial cells were present throughout the peri-implantation window, although relatively much more so in the luminal compared to glandular epithelium. It is conceivable that p16^+^ luminal epithelial cells play a role in directing the embryo to preferential sites of implantation. However, the abundance of p16^+^ cells in both epithelial compartments peaked on the transition of mid- to late-luteal phase, which in turn points towards a potential role for cellular senescence in rendering the endometrium refractory to further implantation.

Clinically, recurrent pregnancy loss (RPL) is a distressing disorders that often remain unexplained despite extensive investigations (Lucas et al., 2016a). Embryonic chromosome instability accounts for a majority of sporadic failures. However, the likelihood of an underlying endometrial defect compromising the development of a euploid embryo increases with each additional failure. Nevertheless, the cumulative live birth rate following multiple miscarriages or IVF failures is high (Lucas et al., 2016a; Smith et al., 2015), which suggests that embryo-endometrial interactions are intrinsically dynamic. Our findings point towards a new paradigm that accounts for the observation that RPL does not preclude a successful pregnancy. If the level of replicative stress during the follicular phase is efficiently counterbalanced by uNK cell mediated clearance of senescence decidual cells during the luteal phase, implantation competence of the endometrium is assured and, in the absence of other pathology, reproductive fitness should be maximal. If not, the frequency of aberrant cycles, and thus the likelihood of reproductive failure, is predicted to increase in line with the degree of endometrial dyshomeostasis. For example, the endometrium in RPL patients is characterized by MSC deficiency, heightened cellular senescence and a prolonged and disordered decidual inflammatory response (Lucas et al., 2016b; Salker et al., 2012). Our model predicts that excessive decidual senescence can be counterbalanced by increased uNK cell proliferation and activation, thus tending towards homeostasis and leading to intermittent normal cycles. Importantly, the degree of endometrial MSC deficiency correlates with the number of previous miscarriages and, by extension, the likelihood of further failure (Lucas et al., 2016b). This observation provides credence to our assertion that the chance of a successful pregnancy correlates inversely with the severity of endometrial dyshomeostasis. The corollary of an intrinsic ability to balance induction and clearance of senescent cells from cycle to cycle is that the human endometrium seems refractory to ageing and maintains its function throughout the reproductive years.

In summary, acute senescence of a subpopulation of stromal cells upon decidualization triggers a multi-step process that transforms the cycling endometrium into a gestational tissue. Endometrial remodeling at the time of embryo implantation is controlled spatiotemporally by the level of decidual senescence and the efficacy of immune clearance.

## Experimental Procedures

### Patient recruitment and sample collection

This study was approved by NHS National Research Ethics Committee (1997/5065). Participants provided written informed consent in accordance with the Declaration of Helsinki, 2000. A total of 2,131 biopsies were used in this study, including 109 samples processed for primary EnSC cultures. Patient demographics are summarized in Table S2. See Supplemental Experimental Procedures for details.

### Decidualization of EnSCs and uNK cell isolation

Primary EnSC cultures were decidualized with 0.5 mM 8-bromo-cAMP and 1 μM medroxyprogesterone acetate (MPA). For co-culture experiments, the supernatant from freshly isolated EnSCs was collected 6-18 h post-seeding and uNK cells isolated using magnetic activated cell separation (MACS; Miltenyi Biotec, Bergisch Gladbach, Germany). See Extended Experimental Procedures for details. In co-culture, the ratio of EnSCs to uNK cells was 2:1. See Supplemental Experimental Procedures for details.

### siRNA transfection

Confluent EnSCs in 24-well plates were transfected using jetPRIME Polyplus transfection reagent (VWR International, Lutterworth, UK) according to the manufacturer’s instructions. Culture medium was refreshed 18 h post-transfection. See Supplemental Experimental Procedures for details.

### Colony-forming unit (CFU) assay

CFU assays were performed as described (Lucas et al., 2016b). Briefly, 500 EnSCs per well were seeded into 10μg/ml fibronectin-coated 6-well plates and cultured in 10% DMEM/F12 containing 10 ng/ml basic fibroblast growth factor for 12 days. Cells were stained with hematoxylin and colonies of more than 50 cells were counted. Cloning efficiency (%) was calculated as the number of colonies formed / number of cells seeded × 100.

### Real-time Quantitative (RTq)-PCR

Total RNA was isolated using STAT-60 (AMS Biotechnology, Oxford, UK), reverse transcribed and subjected to real-time PCR using Power SYBR Green Master Mix (Fisher Scientific, Loughborough, UK), according to manufacturers’ instructions. See Supplemental Experimental Procedures for details.

### Enzyme-linked immunosorbent assay

Detection of individual secreted factors was achieved by commercially available DuoSet ELISA kits (BioTechne, MN, USA) according to the manufacturer’s instructions. See Supplemental Experimental Procedures.

### Senescence-Associated-P-Galatosidase (SAβG)

SAβG staining was performed on confluent EnSC in 24-well plates using Senescence β-Galactosidase Staining Kit (Cell Signalling Technology, MA, USA) according to the manufacturer’s instruction. SAβG activity in cell and tissue lysates was quantified using the 96-Well Cellular Senescence Activity Assay kit (Cell Biolabs Inc; CA, USA). Activity was normalized to protein content, as determined by Bradford assay (Sigma Aldrich, UK). See Supplemental Experimental Procedures for details.

### Western blot analysis

Protein lysates (25 μg per lane) were separated in 12% poly-acrylamide gels by standard SDS-PAGE electrophoresis. Proteins were transferred onto nitrocellulose (GE Healthcare, Amersham, UK) and probed with antibodies targeting Lamin B1 (Abcam, Cambridge, UK; 1:1000), HMGB2 (Abcam; 1:500), p16^INK4^ (Abcam; 1:1500) and p53 (Agilent Technologies, Santa Clara; 1:3000), Histone H1 (Abcam; 1:2000), MacroH2A (Abcam; 1:5000) H3k9me3 (Abcam; 1:1000), FOXO1 (Cell Signaling Technologies, Denvers, M.A, USA; 1:1000) and β-actin (Sigma, Poole; UK), 1:100000). See Supplemental Experimental Procedures for details.

### Immunocytochemistry

Cytospin preparations from 100,000 uNK cells were fixed in 10% formalin and probed with anti-CD56 antibody (Agilent Technologies) (1:250, overnight, 4°C). CD56^+^ cells were identified using the Novolink™ polymer detection system exactly as per manufacturer’s instructions (Leica Biosystems). For immunofluorescence analysis, EnSCs seeded in 35mm glass-bottomed culture dishes were treated and then fixed in 4% paraformaldehyde and permeabilized in 0.5% Triton X-100. See Supplemental Experimental Procedures.

### Immunohistochemistry

Formalin fixed paraffin embedded endometrial sections were stained for CD56 (NCL-L-CD56-504, Novocastra, Leica BioSystems; 1:200) or p16^INK4^ (CINtec^®^ clone E6H4, Roche, Basel, Switzerland; 1: 5). See Supplemental Experimental Procedures for details.

#### RNA-sequencing

Total RNA isolated from snap frozen LH-timed endometrium biopsies from 10 women aged ≤ 30 years and 10 women aged ≥ 40 years was processed for RNA-sequencing. See Supplemental Experimental Procedures.

### Statistical analysis

GraphPad Prism v6 (GraphPad Software Inc.) was used for statistical analyses. Data were checked for normal distribution using Kolmogorov-Smirnov test. Unpaired or paired *t*-test was performed as appropriate to determine statistical significance between two groups. For larger data sets, significance was determined using one-way ANOVA and Tukey’s post-hoc test for multiple comparisons. *P* < 0.05 was considered significant.

## Footnotes

**Author Contributions:** Conceptualization, J.J.B.; Methodology, P.J.B., P.V., E.S.L, S.O and M.H.; Investigation, Y.M., P.J.B., K.F., R.F., J.M., T.Y., L.W., R.L., Y.H.L. and M.H.; Writing – Original Draft, J.J.B., P.J.B and E.S.L.; Funding Acquisition, S.Q., S.T and J.J.B.; Resources, K.F., Sh.T., M.H., S.Q. and J.J.B.; Supervision, M.H., S.Q and J.J.B.

## Acknowledgement

We are grateful to all the women who participated in this research. We thank Gnyaneshwari Patel, Anatoly Shmygol, and Jesús Gil for advice and technical assistance. This work was supported by funds from the Tommy’s National Miscarriage Research Centre and the Biomedical Research Unit in Reproductive Health.

